# Crosstalk between the RNA-binding proteins Regnase-1 and -3 shapes mast cell survival and cytokine expression

**DOI:** 10.1101/2024.01.24.577016

**Authors:** Marian Bataclan, Cristina Leoni, Simone G. Moro, Matteo Pecoraro, Elaine H. Wong, Vigo Heissmeyer, Silvia Monticelli

## Abstract

Post-transcriptional regulation of immune-related transcripts by RNA-binding proteins (RBPs) impacts immune cell responses, including mast cell functionality. Despite their importance in immune regulation, the functional role of most RBPs remains to be understood. By manipulating the expression of specific RBPs in mast cells, coupled with mass spectrometry and transcriptomic analyses, we found that the Regnase family of proteins acts as a potent regulator of mast cell physiology. Specifically, Regnase-1 is required to maintain basic cell proliferation and survival, while both Regnase-1 and −3 cooperatively regulate the expression of inflammatory transcripts upon mast cell activation, with *Tnf* being a primary target of both proteins. In mast cells, Regnase-3 directly interacts with Regnase-1 and is necessary to restrain Regnase-1 expression through the destabilization of its transcript. Overall, our study identifies protein interactors of endogenously expressed Regnase factors, characterizes the regulatory interplay between Regnase family members in mast cells, and establishes their role in the control of mast cell homeostasis and inflammatory responses.

## Introduction

Mast cells are tissue-resident immune cells that exert key effector functions during allergic and anaphylactic reactions, act as sentinels against invading pathogens, and regulate physiological processes, such as tissue repair and angiogenesis (1, 2). These cells are activated by a variety of signals through multiple receptors, most commonly via the high-affinity IgE receptor, FcεRI. Recognition of antigens or allergens by IgE antibodies bound to FcεRI triggers cell activation, characterized by degranulation and rapid release of a broad panel of pre-stored and *de novo* synthesized mediators (histamine, prostaglandin, cytokines, proteases, etc.), eventually resulting in an inflammatory response (2, 3). Dysregulation of mast cell activation and functions may lead to excessive and damaging responses to otherwise harmless agents, as exhibited in mast cell activating syndromes and IgE-associated disorders (food allergy, allergic rhinitis, asthma and anaphylaxis) (2). Tight control of mast cell functionality is therefore crucial to facilitate responses that will effectively eradicate a pathogen or allergen, while minimizing harm to the host.

Post-transcriptional regulation of immune-related mRNAs is critical in controlling immune cell responses. This can be mediated by RNA-binding proteins (RBPs), which bind to specific elements within the RNA transcripts and regulate different post-transcriptional events, such as RNA processing, modifications, stability, and translation (4–6). Mechanistically, RBPs can restrain inflammation by binding to *cis*-regulatory regions, such as AU-rich regions or stem-loop structures, in the 3’ untranslated region (UTR) of inflammatory mRNAs, leading to the destabilization and eventual degradation of these transcripts. In mast cells, we recently showed that m^6^A mRNA methylation is crucial to restrain mast cell responses, and that m^6^A methylation of the *Il13* transcript determines its stability and the extent of IL-13 cytokine production (7). However, the mechanistic details regarding the functions of many RBPs and their role in the regulation of mast cell responses are yet to be elucidated.

Here, we found that the expression of members of the Regnase family of RBPs is highly induced upon acute stimulation of mast cells. Regnase proteins (Regnase-1 to −4, encoded by the *Zc3h12a-d* genes) are essential to restrain excessive inflammatory responses in immune cells, through the action of an intrinsic RNase enzymatic activity (8, 9). Upon recognition of stem-loop structures in target mRNAs, Regnases directly cleave and destabilize inflammatory transcripts (8). Linked to such prominent role in restraining effector immune responses, Regnase-1 has recently gained attention as a candidate therapeutic target to enhance anti-tumor responses of CD8^+^ T lymphocytes. Deletion of Regnase-1 was sufficient to improve T cell expansion, functionality and persistence, leading to enhanced anti-tumor responses in mice (10–13). On the other hand, significant amelioration of disease was also observed in preclinical models of autoimmunity in which the expression of Regnase-1 was stabilized by the *in vivo* injection of morpholino oligonucleotides. These oligonucleotides effectively blocked the stem-loop structures in the *Zc3h12a* 3’ UTR that are normally targeted by Regnase-1 itself in a process of autoregulation, resulting in the observed amelioration of the disease (14). Within the myeloid compartment, Regnase-1 was shown to prominently regulate a number of inflammatory transcripts in macrophages, including *Il6, Il12b* and *Ptgs2* (9, 15, 16). Differently from Regnase-1, which is expressed both in lymphoid and myeloid cells, Regnase-3 functions primarily in myeloid cells, and its macrophage-specific deletion led to excessive IFN-γ expression (17). Despite the growing interest in understanding the role of Regnases in immune response regulation, numerous questions persist regarding their mechanism of action. These include the extent of their unique or redundant functions in immune cells, and whether they physically interact to modulate shared mRNA targets.

By leveraging a variety of experimental approaches, such as gene editing and genetic deletions of Regnase proteins, modulating their expression via mRNA delivery, and identifying their protein interactome through immunoprecipitation coupled with mass-spectrometry, our study reveals that Regnase-1 serves as a key negative regulator of mast cell proliferation and survival. In contrast, Regnase-3 primarily modulates the expression of Regnase-1 within mast cells, with which it also physically interacts. Despite showing some distinct regulatory roles, influenced in part by the preference of these two proteins for different subcellular compartments, both Regnase-1 and −3 share regulatory control over a subset of inflammatory transcripts, including *Tnf*. In summary, our study delineates the protein interactome of endogenously expressed Regnase proteins and elucidates their significance in governing mast cell homeostatic and inflammatory responses.

## Results

### Expression of both Regnase-1 and −3 is induced in acutely stimulated mast cells

To determine how the expression of RBP genes changes in mast cells upon stimulation with IgE and antigen complex, we first re-analyzed RNA-seq datasets from Li et al. (18) focusing only on RBP genes as defined in the RBP2go database (19) and including only RBPs detected in more than one dataset (**Suppl. Figure 1a** and **Suppl. Table 1**). This resulted in overall 827 differentially expressed transcripts, of which 435 were associated with general RNA metabolic processes at gene ontology (GO) analysis. Within this category, RBP transcripts divided into further functional categories linked primarily to RNA processing, translation and stability (**Suppl. Figure 1b**). Next, we found that *Zc3h12c* (Regnase-3) was among the most highly induced RBP-encoding transcripts following stimulation (**Suppl. Figure 1a**). Two other Regnase family members, *Zc3h12a* (Regnase-1) and *Zc3h12d* (Regnase-4) were also induced, while *Zc3h12b* (Regnase-2) was downmodulated. Re-analysis of ATAC-seq data from Li et al. (18) also showed increased accessibility around the *Zc3h12a* and *Zc3h12c* promoters upon IgE crosslinking, suggesting transcriptional activation after acute stimulation (**Suppl. Figure 1c**). Finally, analysis of the expression of *Zc3h12a-d* in mast cells extracted *ex-vivo* from different mouse tissues (ImmGen database) (20), also revealed high expression of *Zc3h12c* in most tissue mast cells (notably in skin mast cells, less prominently in peritoneal cavity mast cells (PMCs)), followed by modest expression of *Zc3h12a* and very low levels of *Zc3h12b* and *Zc3h12d* (**Suppl. Figure 1d**). Overall, these observations point towards a role for selected Regnase family members in modulating mast cell functions, and we therefore focused our attention on this family of RBPs.

To validate and extend these findings, we first measured the expression of the Regnase-encoding transcripts in bone marrow-derived mast cells (BMMCs, ∼99% pure populations, determined by FcεRIα and c-Kit staining, **Suppl. Figure 1e**) activated with IgE and antigen complexes for the indicated times. *Zc3h12b* and *Zc3h12d* were expressed at very low levels and were not significantly induced upon stimulation, while both *Zc3h12a* and *Zc3h12c* were expressed and highly induced at early time points (within the first 1-2h) in activated mast cells (**Figure 1a**). Similar significant induction specifically of *Zc3h12a* and *Zc3h12c* was observed in peritoneal mast cells separated *ex-vivo* from the peritoneal cavity and stimulated with PMA and ionomycin (**Figure 1b** and **Suppl. Figure 1f**). Regnase-1 and −3, were also induced at the protein level upon activation of BMMCs and peritoneal cavity-derived, *in-vitro* expanded mast cells (PCMCs) (**Figure 1c-d**), pointing towards a role for these two Regnase family members in regulating mast cell responses to stimuli. We also detected the expected proteolytic cleavage of Regnase-1 upon activation, consistent with Malt-1 activation downstream of the IgE receptor complex (8, 21). Whether Regnase-3 is regulated post-translationally remains to be understood, especially considering the complex pattern revealed by western blot (**Figure 1c-d**), as reported also in the literature (17). In terms of subcellular localization, both Regnase-1 and −3 were detected within punctuated structures in the cytoplasm of acutely activated mast cells, with limited overlap (**Suppl. Figure 2**). Because mast cells can be activated by a large variety of signals, we determined whether stimuli other than IgE and antigen complexes affected Regnase expression. We found that both *Zc3h12a* and *Zc3h12c* were induced, albeit at different extent, by inflammatory signals that induce cytokine expression in mast cells, including IL-33 and LPS, but not by the c-Kit ligand stem cell factor (SCF), which by itself did not induce cytokine expression (**Figure 1e** and **Suppl. Figure 3**). Overall, these data point towards a role for Regnase-1 and − 3 in modulating mast cell activation and/ or effector functions, that we set out to investigate.

**Figure 1.**
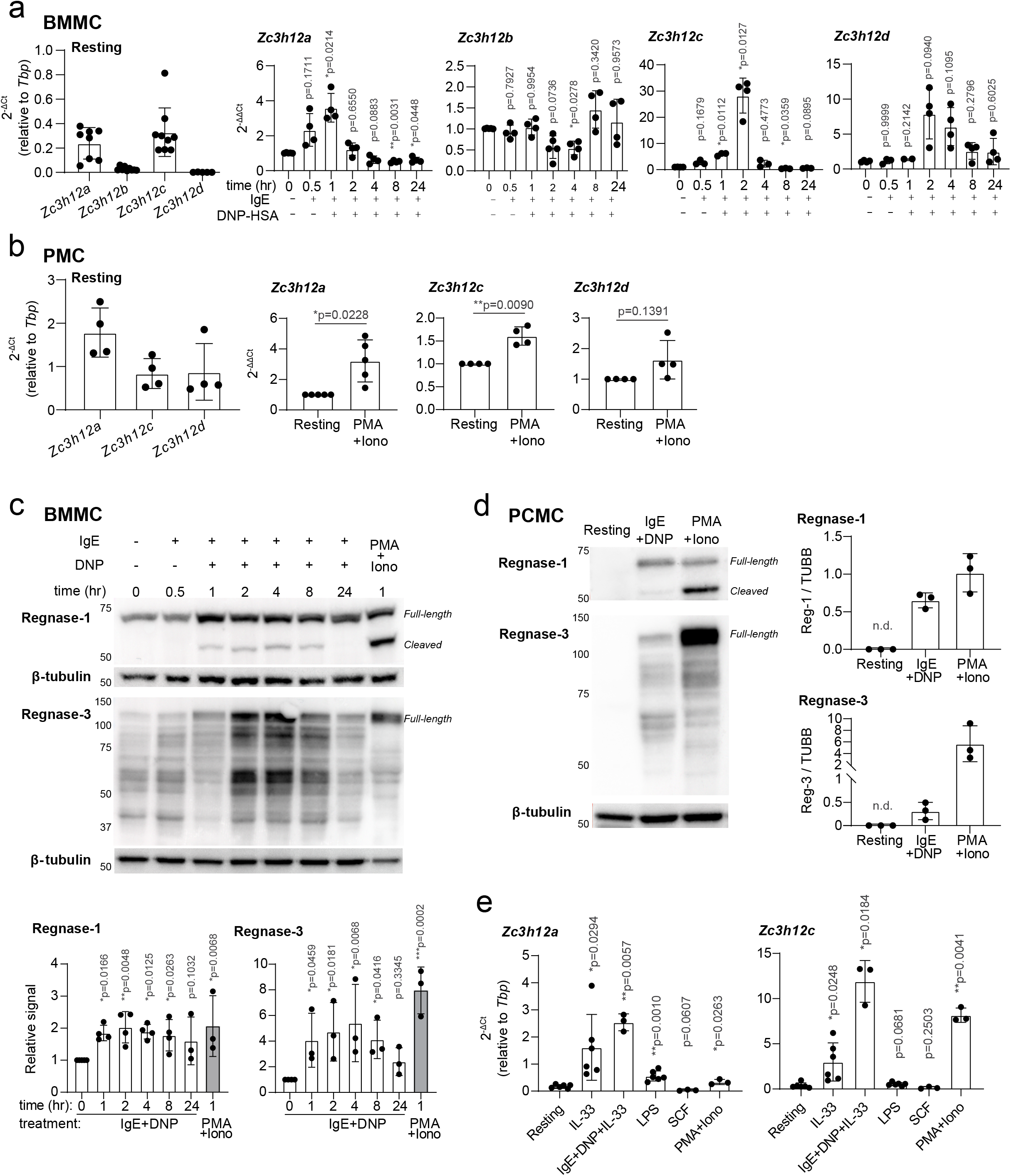
Expression of Regnase-1 and Regnase-3 is induced upon mast cell activation. **a)** Expression of *Zc3h12a-d* in BMMCs at resting state (left panel) and upon activation with IgE and antigen complexes for up to 24 h (right), normalized to *Tbp* expression. N=4 to 9 independent experiments. Mean ± SD. One-way ANOVA. **b)** Expression of *Zc3h12a-d* in *ex vivo*-isolated PMCs at resting state (left) and upon activation with PMA and ionomycin for 2 h (right), normalized to *Tbp* expression. N=4 to 5 independent experiments. Mean ± SD. Paired t-test, two-tailed. **c)** Expression of Regnase-1 and Regnase-3 in BMMCs activated with IgE and antigen complexes or PMA and ionomycin at various time points. Regnase-1 (full-length + cleaved) and Regnase-3 (full-length) levels were normalized to β-tubulin expression. N=3 to 4 independent experiments. Mean ± SD. One-way ANOVA. **d)** Expression of Regnase-1 and Regnase-3 in cultured PCMCs activated with IgE and antigen complexes or PMA and ionomycin for 2 h, normalized to β-tubulin expression. N=3 independent experiments. Mean ± SD. n.d.: not determined. **e)** Expression of *Zc3h12a* and *Zc3h12c* upon activation of BMMCs with different stimuli for 2 h, normalized to *Tbp* expression. N=3 to 6 independent experiments. Mean ± SD. Paired t-test, two-tailed.

### Regnase-1 and −3 restrain the expression of inflammatory cytokines in mast cells

To gain a broad understanding of the impact of Regnase-1 and −3 on inflammatory transcripts, we first used siRNAs to deplete both factors (schematics of siRNA targeting strategy in **Figure 2a**). Depletion of both Regnase-1 and −3, alone and in combination, was highly effective at the mRNA and protein level (**Figure 2b-c**). Notably, depletion of Regnase-3 led to significantly increased expression of Regnase-1, pointing towards a role for Regnase-3 in restraining Regnase-1 expression (**Figure 2c**), consistent with previous data described in macrophages (17, 22). Next, we stimulated the cells for 2 h with IgE and antigen, and we measured the expression of 734 immune transcripts by Nanostring digital profiling. We found that 325 of these transcripts were expressed by mast cells, and that the depletion of Regnase-1, alone or in combination with Regnase-3, led to increased expression of mRNAs encoding inflammatory mediators, including established Regnase-1 targets such as *Nfkbiz* and *Il2* (8, 23) (**Figure 2d** and **Suppl. Table 2**). Differently from Regnase-1, depletion of Regnase-3 alone had only a very modest impact on the inflammatory transcriptome of mast cells, with only 9 genes significantly upregulated (log_2_ fold-change >0.5 and p-value ≤ 0.05) (**Figure 2d**). Depletion of both proteins at the same time led to an overall combined phenotype that enhanced the effect of the individual depletions, with transcripts such as *Tnf, Il1b*, *Il2* and *Nfkbiz* becoming even more prominently dysregulated (**Figure 2d**), suggesting redundant functions at least on specific targets. Overall, we found that Regnase-1, and to a lesser extent Regnase-3, restrain the expression of inflammatory transcripts in mast cells. The downregulated transcripts were likely to be the result of indirect effects of Regnase depletion, and no previously described (9) Regnase-bound, direct targets were identified among the downregulated transcripts (**Figure 2e**). Vice versa, among the significant upregulated genes several direct Regnase-1 targets could be observed (red asterisks, **Figure 2e**), suggesting that our analysis identified transcripts at least in part directly modulated by Regnase proteins. Interestingly, *Il2* and *Tnf* were among the few inflammatory transcripts significantly increased upon depletion of both Regnase-1 and −3, alone or in combination. Since *Il2* was expressed at low levels in mast cells, we further investigated the impact of Regnases on TNF expression. First, we confirmed increased expression of the *Tnf* transcript in an independent set of experiments (**Figure 3a**). Next, we found that, concordant with the mRNA data, TNF protein expression was significantly increased upon depletion of Regnase-3, alone or in combination with Regnase-1, while the impact of Regnase-1 depletion alone was more modest (**Figure 3b**). Next, to confirm these findings using an independent experimental setup, we used CRISPR-Cas9-mediated gene editing to deplete the expression of *Zc3h12a* or *Zc3h12c* (**Figure 3c**). Depletion of either protein was highly effective in mast cells, as shown by western blot (**Figure 3d**). Concordant with our siRNA data, depletion of both Regnase-1 and −3 led to significantly increased *Tnf* mRNA expression (**Figure 3e**), as well as to increased protein expression, measured by intracellular staining, both in response to IgE and antigen complexes or PMA and ionomycin stimulation (**Figure 3f** and **Suppl. Figure 4a**). We also measured the expression of other cytokines prominently produced by mast cells. We found that IL-6 expression was modestly affected by Regnase-1 deletion (**Suppl. Figure 4a**), consistent with our Nanostring data, showing mildly (and not significant) increased expression, and it was not affected by Regnase-3 deletion. Although at the mRNA level we could not detect any effect on the *Il13* transcript, IL-13 expression was significantly increased upon deletion of either Regnase, suggesting indirect regulation.

**Figure 2.**
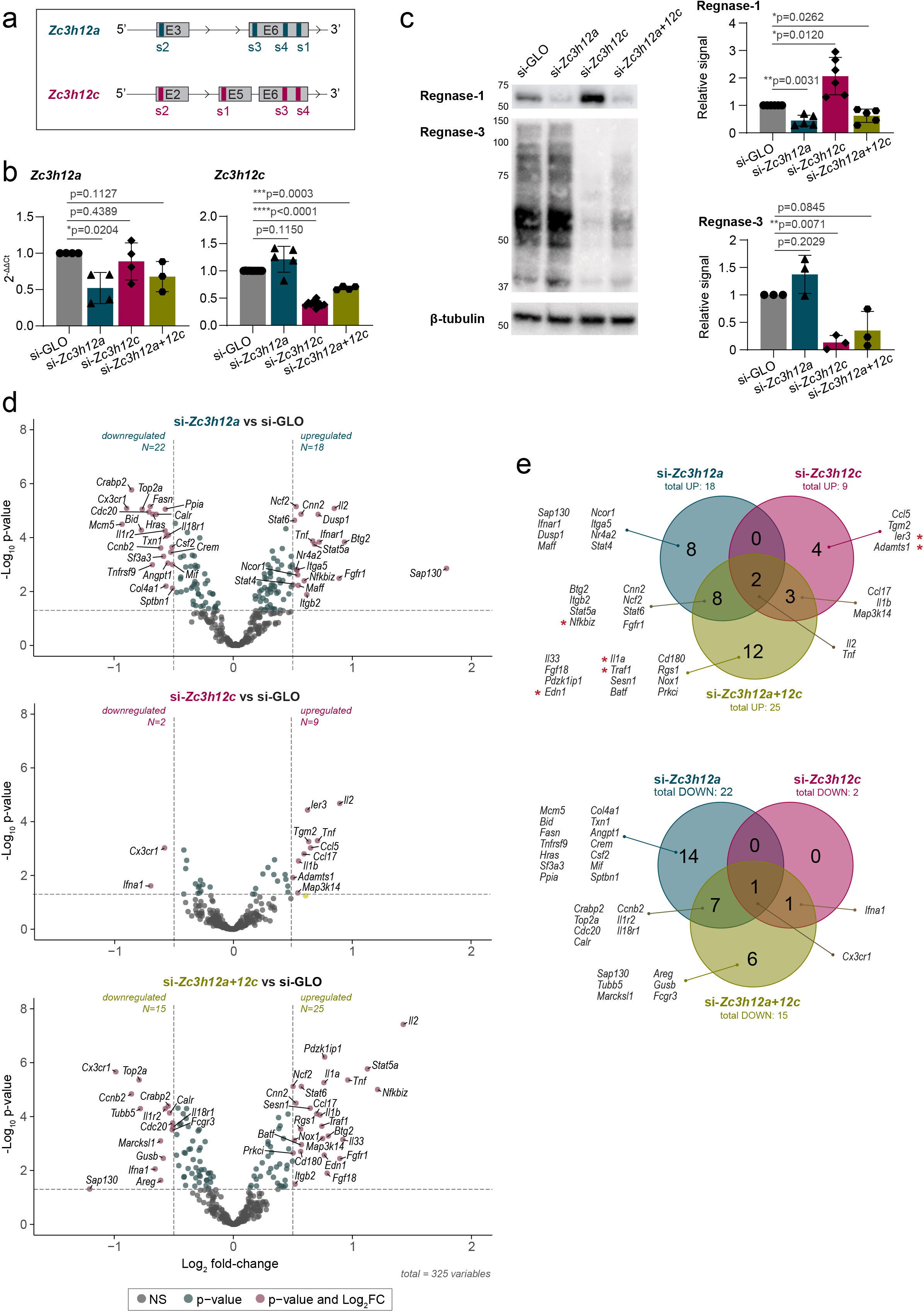
Loss of Regnase-1 and Regnase-3 induces inflammatory gene expression in activated mast cells. **a)** Schematic representation of *Zc3h12a* and *Zc3h12c* loci with regions targeted by siRNAs for RNAi-mediated gene knockdown. **b-c)** mRNA **(b)** and protein **(c)** expression of Regnase-1 and Regnase-3 upon transfection of BMMCs with siRNAs targeting *Zc3h12a* and/or *Zc3h12c*. mRNA expression is normalized to *Tbp*, while protein expression is normalized to β-tubulin. N=3 to 6 independent experiments. Mean ± SD. Paired t-test, two-tailed. **d)** Differentially expressed genes (−0.5>log_2_ fold-change>0.5) in Regnase-1 and/or Regnase-3 knockdown BMMCs activated with IgE and antigen complexes for 2 h, measured by Nanostring digital profiling. **e)** Venn diagram of differentially expressed genes in **(d)**. Genes marked with a red asterisk were reported to be directly bound by Regnase-1 in a RIP-seq analysis of HeLa cells (9).

**Figure 3.**
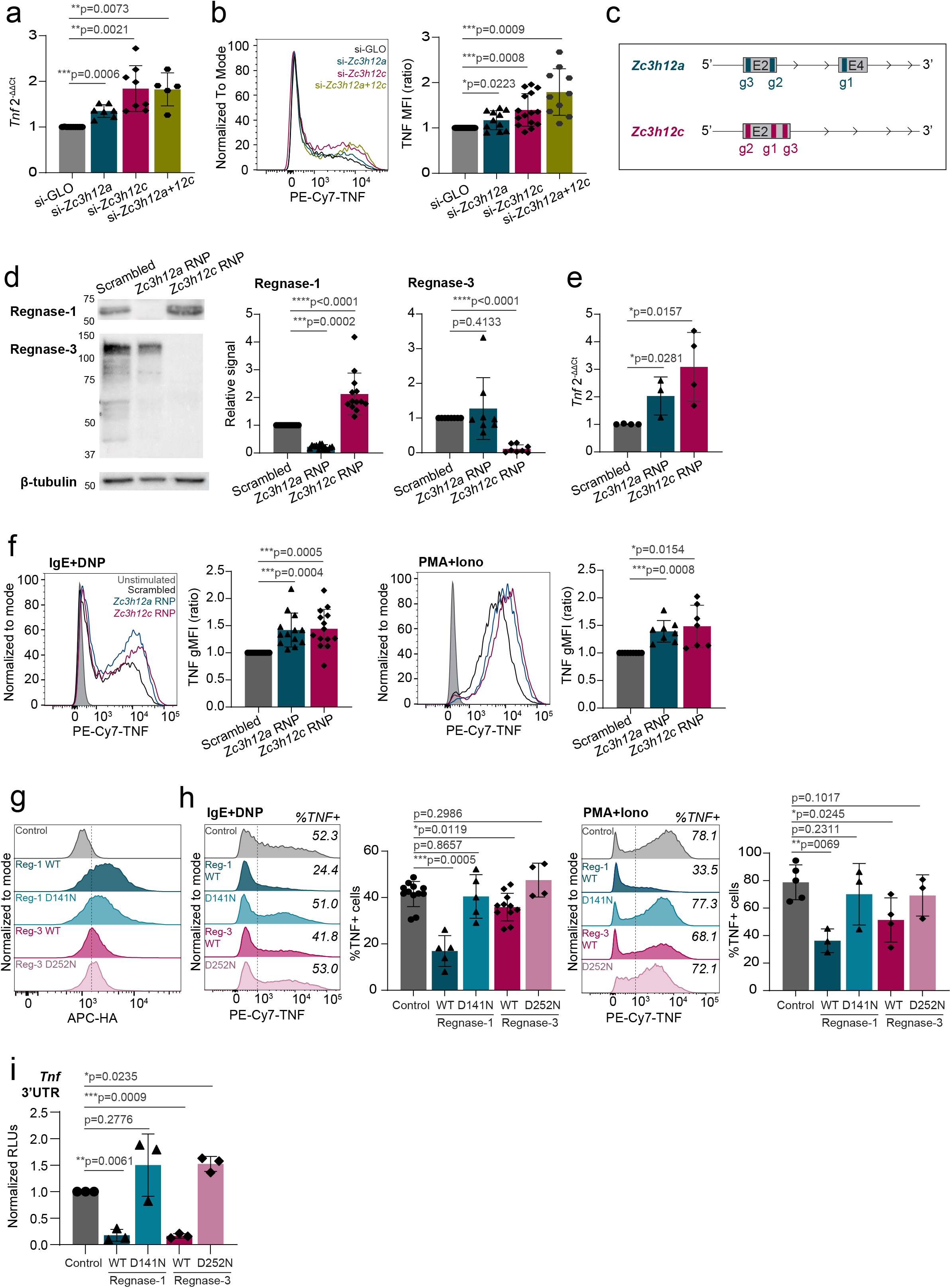
Regnase-1 and Regnase-3 negatively regulate TNF expression. **a-b)** mRNA expression **(a)** and intracellular staining **(b)** of TNF in activated BMMCs (IgE and antigen complexes for 2 h in **(a)** and 4 h in **(b)**) transfected with siRNAs against *Zc3h12a* and/or *Zc3h12c*. mRNA expression is normalized to *Tbp*. Representative histogram and mean fluorescence intensity (normalized to siGLO) of TNF are shown. N=5 to 13 independent experiments. Mean ± SD. Paired t-test, two-tailed. **c)** Schematic representation of *Zc3h12a* and *Zc3h12c* loci with regions targeted by guide RNAs (gRNAs) for CRISPR-Cas9-mediated gene knockout. **d)** Expression of Regnase-1 and Regnase-3 upon transfection of BMMCs with CRISPR-Cas9-gRNA RNPs targeting *Zc3h12a* or *Zc3h12c* after 1 week of culture, normalized to β-tubulin expression. N=8 to 13 independent experiments. Mean ± SD. Paired t-test, two-tailed. **e)** *Tnf* expression in Regnase-1 and Regnase-3 knockout BMMCs activated with IgE and antigen complexes for 2 h, normalized to *Tbp* expression. N=3 to 4 independent experiments. Mean ± SD. Unpaired t-test, two-tailed. **f)** Intracellular staining of TNF in Regnase-1 and −3 knockout cells activated with IgE and antigen complexes (left) or PMA and ionomycin (right) for 4 h. N=8 to 14 independent experiments. Mean ± SD. Paired t-test, two-tailed. **g)** Intracellular staining of HA-tag in BMMCs transfected with *in vitro-* transcribed HA-tagged *Zc3h12a* or *Zc3h12c* mRNA to assess overexpression of Regnase-1 and −3, either wild-type or RNase-inactive mutant (D141N or D252N). **h)** Intracellular staining of TNF in Regnase-1- and −3-overexpressing cells activated with IgE and antigen complexes (left) or PMA and ionomycin (right) for 4 h. N=3 to 11 independent experiments. Mean ± SD. Paired t-test, two-tailed. **i)** Luciferase reporter assay upon co-transfection of Regnase-1 or −3, either wild-type or RNase-inactive mutant (D141N and D252N), with *Tnf* 3’UTR reporter plasmid. N=3 independent experiments. Mean ± SD. Paired t-test, two-tailed.

To confirm these data, we sought to force expression of Regnase enzymes in mast cells. However, we found that most overexpression methods, including inducible vectors, failed to provide stable Regnase-1 and −3 protein expression, due to deleterious effects especially of Regnase-1 on mast cell viability. We therefore optimized an *in vitro* transcription (IVT) system for the direct, short-term delivery of mRNAs encoding Regnase-1 and −3. The T7-driven transcripts were stabilized by the addition of the 5’- and 3’-UTRs from the *HBB* gene, followed by 5’-capping and 3’-poly(A) tailing. We generated transcripts leading to the expression of HA-tagged, wild-type Regnase-1 and −3 and versions carrying point mutations in the RNase enzymatic domain (D141N and D252N, respectively), leading to disruption of the catalytic activity (16). mRNA delivery of Regnase-1 and −3, either wild-type or RNase mutant, proved to be feasible and efficient, and all proteins were highly expressed in transfected mast cells, as measured by intracellular staining using an antibody against the HA-tag (**Figure 3g** and **Suppl. Figure 4b**). Cell viability was comparable across conditions (**Suppl. Figure 4b**), and the overall subcellular localization of the IVT mRNA-derived proteins was comparable to that of endogenously expressed Regnase-1 and −3 (**Suppl. Figure 5**). Analysis of cytokine expression by these cells upon stimulation with either IgE and antigen complexes or PMA and ionomycin, revealed that both Regnase-1 and −3 significantly diminished the ability of mast cells to produce TNF in an RNase-dependent manner (**Figure 3h**). Compared to Regnase-1, the somewhat reduced capability of Regnase-3 to limit TNF expression was likely linked to the extent of overexpression that we could achieve in this experimental system, that was less pronounced for Regnase-3 (**Figure 3g**). Mostly consistent with our CRISPR-Cas9 knock-out data, overexpression of Regnase-3 had no detectable effect on IL-6 production, while Regnase-1 limited both IL-6 and IL-13 expression by mast cells (**Suppl. Figure 6**). To determine whether the effect of Regnase-1 and −3 on TNF expression was mediated by the direct targeting of the *Tnf* transcript, we cloned the full-length 3’UTR of *Tnf* into a luciferase reporter vector. We then co-transfected it with Regnase-1 or Regnase-3-expressing vectors, either wild-type or RNase-impaired. We found that the *Tnf* 3’UTR was strongly affected by both Regnases, and that luciferase expression was restored upon mutation of their RNase enzymatic sites, pointing towards a direct effect on *Tnf* that required the RNase enzymatic activity of Regnase proteins (**Figure 3i**). Overall, both Regnase-1 and −3 are required to effectively limit TNF expression by mast cells, although their effect on other inflammatory cytokines appears to be more selective and predominantly linked to Regnase-1.

### Regnase-3 is required to restrain expression of Regnase-1 in mast cells

Depletion of Regnase-3 by RNAi led to significantly increased Regnase-1 expression in mast cells (**Figure 2c**), indicating that understanding the phenotype of either knock-down or knock-out is complicated by the presence of direct and indirect effects and cross-regulation between these two factors. To better dissect the role of Regnase-3 in regulating Regnase-1 expression, we first confirmed the effect of Regnase-3 on the *Zc3h12a* mRNA by RNAi in an independent set of experiments, and by deleting *Zc3h12c* by CRISPR-Cas9. In both experimental setups, *Zc3h12a* mRNA expression was significantly increased (**Figure 4a**). Concordant with this result, we found that upon overexpression of Regnase-3 by IVT mRNA transfection, Regnase-1 protein expression was significantly reduced (**Figure 4b**). Regnase-1 is also known to diminish the stability of its own transcript through stem-loop structures in the 3’UTR (14, 15). To assess the impact of the individual Regnases on *Zc3h12a* expression, we cloned the 3’UTR of *Zc3h12a* in a luciferase reporter plasmid and measured its response upon co-transfection of Regnase-1 or −3. Both proteins were able to strongly affect luciferase expression in an RNase activity-dependent manner, suggesting a direct effect on the 3’UTR of *Zc3h12a* (**Figure 4c**). Finally, to determine whether Regnase-3 was required to modulate *Zc3h12a* transcript stability in mast cells, we treated the cells with actinomycin D, followed by the measurement of *Zc3h12a* mRNA levels over time. Deletion of Regnase-3 was sufficient to increase the half-life of the *Zc3h12a* transcript in mast cells (**Figure 4d**). On the contrary, we found no evidence of any impact of Regnase-1 depletion or deletion on Regnase-3 (or other Regnases) expression, except for a modest upregulation of *Zc3h12b* in absence of Regnase-1 (**Figure 4e-f**).

**Figure 4.**
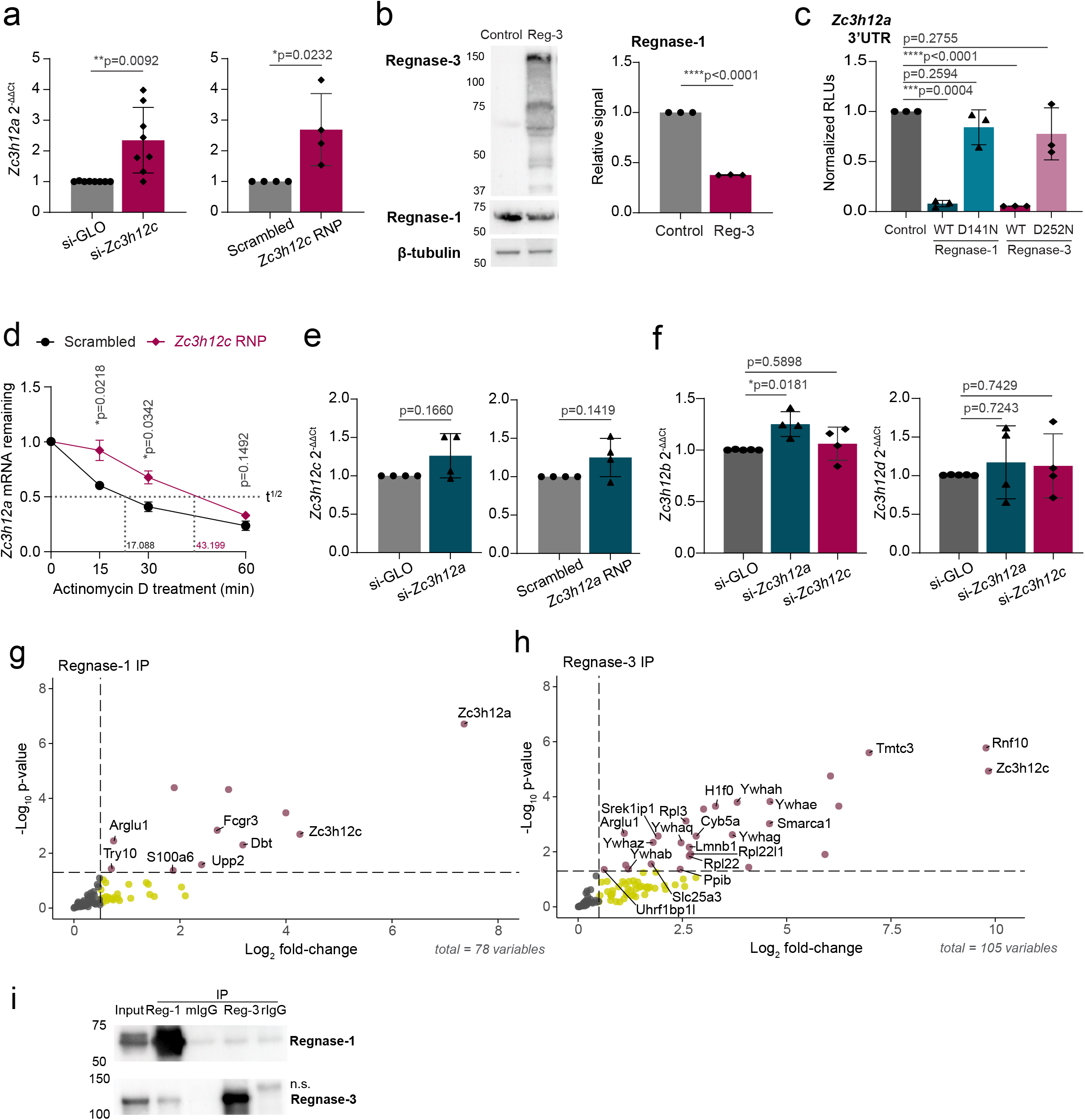
Regnase-3 acts as a negative regulator of Regnase-1 by destabilizing the *Zc3h12a* transcript. **a)** Expression of *Zc3h12a* in Regnase-3 knockdown (left) or knockout (right) BMMCs activated with IgE and antigen complexes for 2 h, normalized to *Tbp*. N=4 to 8 independent experiments. Mean ± SD. Ratio paired t-test, two-tailed. **b)** BMMCs were transfected with *in vitro* transcribed *Zc3h12c* mRNA, followed by western blot to assess the effect of Regnase-3 overexpression on Regnase-1 expression, normalized to β-tubulin. N=3 independent experiments. Mean ± SD. Paired t-test, two-tailed. **c)** HEK293T cells were transfected with plasmids expressing Regnase-1 or −3, either wild-type or RNase-inactive mutant (D141N and D252N), together with a *Zc3h12a* 3’UTR reporter plasmid, followed by luciferase assay. N=3 independent experiments. Mean ± SD. Paired t-test, two-tailed. **d)** Mast cells were transfected with *Zc3h12c* RNPs (or scrambled RNP control), followed by actinomycin D treatment to block transcription. *Zc3h12a* transcript stability was measured by RT-qPCR at the indicated time-points after stimulation with IgE and antigen complexes. N=4 independent experiments. Mean ± SEM. Paired t-test, two-tailed. Half-life (t^1/2^) was calculated using non-linear regression analysis. **e)** Expression of *Zc3h12c* in Regnase-1 knockdown (left) or knockout (right) BMMCs activated for 2 h with IgE and antigen complexes, normalized to *Tbp*. N=4 independent experiments. Mean ± SD. Ratio paired t-test, two-tailed. **f)** Expression of *Zc3h12b* (left) or *Zc3h12d* (right) in Regnase-1 or Regnase-3 knockdown BMMCs activated for 2 h with IgE and antigen complexes, normalized to *Tbp*. N=4 independent experiments. Mean ± SD. Ratio paired t-test, two-tailed. **g-h)** Immunoprecipitation and mass spectrometry analysis of endogenous **(g)** Regnase-1 and **(h)** Regnase-3 in BMMCs activated with PMA and ionomycin for 1 h. Immunoprecipitation with mouse or rat IgG isotype control was used to normalize mass spectrometry data, respectively. Proteins with log_2_FC>0 from 3 to 4 independent experiments are shown. **i)** Western blot analysis upon immunoprecipitation of Regnase-1, Regnase-3, and corresponding isotype control in HEK293T cells co-transfected with Regnase-1 and Regnase-3 for 48 h. mIgG: mouse IgG, rIgG: rat IgG, n.s.: non-specific.

To gain more insights into the cross-talk between Regnase-1 and −3, we performed immunoprecipitation and mass-spectrometry analysis of the endogenously expressed proteins in mast cells activated with PMA and ionomycin (**Figure 4g-h, Suppl. Figure 7a** and **Suppl. Table 3**). We found that immunoprecipitation of Regnase-1 recovered Regnase-3 as its primary endogenous interactor. This interaction was confirmed in HEK293T cells upon co-transfection of Regnase-1 and Regnase-3 (**Figure 4i** and **Suppl. Figure 7b**). On the other hand, Regnase-3 interacted primarily with ribosomal proteins, suggesting an involvement in translation, as well as with the 14-3-3 family of proteins. 14-3-3 proteins were previously shown to interact with FLAG-HA-tagged Regnase-1 when transfected into HeLa cells and to contribute to Regnase-1 protein stability (25). Since in that system Regnase-3 was not expressed, it could not be recovered as an interactor of Regnase-1. However, in our experimental setting in which both Regnase family members are expressed, complex formation involves primarily Regnase-1 and −3 on the one side and Regnase-3 and 14-3-3 proteins on the other, although cell- and stimulus-specific differences may also apply. The fact that in mast cells Regnase-3 was identified as an interactor of Regnase-1, but not the opposite, suggests that Regnase-3 is predominantly interacting with other protein partners, and only a relatively small proportion remains available for Regnase-1 interaction. Overall, our data revealed that Regnase-3 is a crucial direct interactor and regulator of Regnase-1 expression in mast cells.

### Regnase-1, but not Regnase-3, is required to maintain mast cell proliferation and viability

Having established the role of Regnase-1 and −3 in regulating inflammatory responses in mast cells, we investigated their role in modulating mast cell growth and survival. First, depletion of Regnase-1, but not Regnase-3, was sufficient to reduce cell viability 72 h after siRNA transfection, measured by Live/Dead staining (**Figure 5a**). Consistent with this observation, measuring the extent of apoptosis by Annexin V staining revealed a significant increase in the percentage of early apoptotic cells upon Regnase-1 depletion (**Figure 5b**), as well as diminished ability of the cells to proliferate, measured by BrdU incorporation (**Figure 5c**). Cell cycle analysis confirmed an increased percentage of cells in G_1_ and a concomitant decrease in the percentage of cells in S phase (**Figure 5d**). A similar increase in cell death was observed upon deletion (by CRISPR-Cas9) of *Zc3h12a*, that remained significant regardless of the presence of IL-3 (an essential survival factor for mast cells) in the culture medium (**Figure 5e** and **Suppl. Figure 8a**) and was accompanied by reduced overall proliferation (**Figure 5f** and **Suppl. Figure 8b**). Finally, a modest decrease in the capacity of the cells to degranulate in response to IgE and antigen stimulation was observed upon depletion or deletion of Regnase-1 and was likely linked to the disrupted basic cellular functionalities (**Figure 5g**).

**Figure 5.**
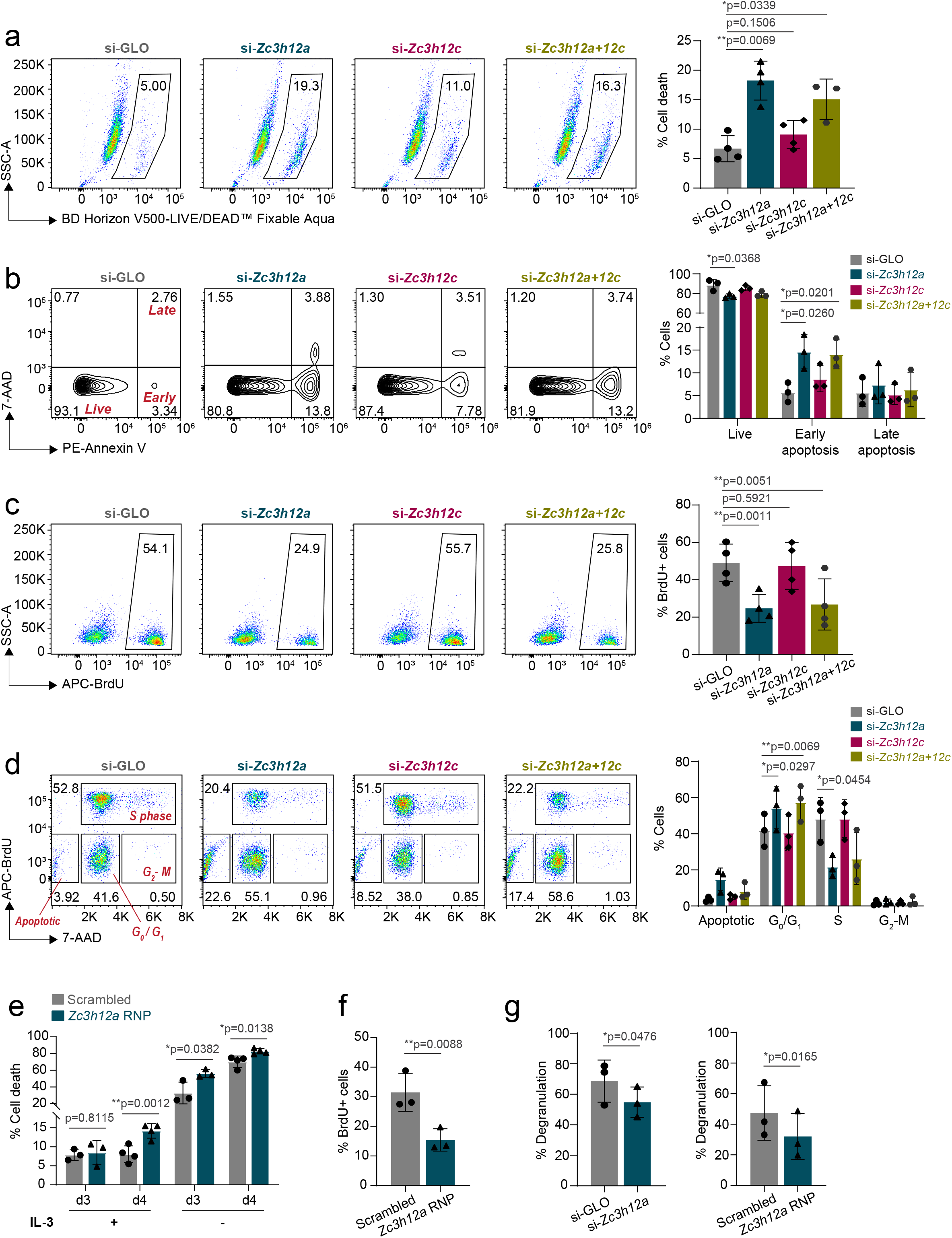
Regnase-1 is crucial in maintaining mast cell homeostatic functions. **a-d)** BMMCs were transfected with siRNAs targeting the indicated Regnase family members, followed by phenotypic analyses 72 h after transfection. **a)** LIVE/DEAD staining to measure cell viability. N=3 to 4 independent experiments. Mean ± SD. Paired t-test, two-tailed. **b)** Annexin V and 7-AAD staining to measure cell apoptosis. N=3 independent experiments. Mean ± SD. Unpaired t-test, two-tailed. **c)** BrdU staining to measure cell proliferation. N=4 independent experiments. Mean ± SD. Paired t-test, two-tailed. **d)** BrdU and 7-AAD co-staining to measure cell cycle progression. N=3 independent experiments. Mean ± SD. Two-way ANOVA. **e-f)** BMMCs were transfected with Cas9-gRNA RNPs targeting *Zc3h12a*. The following experiments were performed after one week: **e)** LIVE/DEAD staining to measure cell viability after growing cells in medium with or without IL-3 for 72-96 hours. N=3 to 4 independent experiments. Mean ± SD. Paired t-test, two-tailed. **f)** BrdU staining to measure cell proliferation. N=3 independent experiments. Mean ± SD. Paired t-test. **g)** Degranulation assay was performed on cells transfected with *Zc3h12a-*targeting siRNAs (left) or Cas9-gRNA RNPs (right) by measuring the extent of β-hexosaminidase release upon cell stimulation with IgE and antigen complexes for 1 h. N=3 independent experiments. Mean ± SD. Ratio paired t-test, two-tailed.

To gain insights into the mechanism(s) that may lead to such pronounced impairment in the ability of mast cells to survive in absence of Regnase-1, we deleted Regnase-1 by CRISPR-Cas9 and performed RNA-sequencing. We found that 131 genes were upregulated while 187 genes were downregulated in absence of Regnase-1, pointing towards both direct and indirect effects (**Figure 6a** and **Suppl. Table 4**). Among the upregulated genes, basal expression of *Tnf* was increased, even in absence of stimulation. Some previously reported direct targets of Regnase-1 (9), including *Nfkbid* and *Ran*, were also modestly upregulated (**Suppl. Table 4**). Transcripts encoding the mast cell protease Mcpt2 and the metalloprotease Adam8 were increased, while some genes involved in regulating mast cell degranulation such as *Hdc* (26) were downregulated. GO analysis of the downregulated genes clustered around signal transduction as well as cholesterol and lipid metabolism. These genes include *Scd1*, *Scd2*, *Acsl3*, and *Fads2*, which are important for lipid synthesis, as well as *Abcg1, Abca1*, *Pcsk9* and *Fabp5,* which are involved in lipid transport, pointing towards overall dysregulation in lipid homeostasis. Metabolism-related genes downmodulated in the absence of Regnase-1 also included *Aldoc*, *Pdk1* and *Hk2,* which play a role in glucose metabolism. These results point towards an involvement of Regnase-1 in regulating metabolic processes required for cell maintenance, as observed also in other cell types (24, 27–29). On the other hand, categories associated to the upregulated genes revealed many GO terms associated to the control of cell cycle, DNA replication and repair (**Figure 6b**). Indeed, some of the most differentially expressed genes included factors important for cell cycle progression, such as *Cdca7*, *Cks1b* and *Cdk1.* While classical anti- and pro-apoptosis-related genes, such as *Bcl2*, *Bax* and *Bcl2l1*, were unchanged, DNA damage and replication markers such as *Aunip, Ung, Pold1,* and *H2ax* were upregulated, suggesting that Regnase-1 knockout cells were more susceptible to cellular stress, in addition to cell cycle defects.

**Figure 6.**
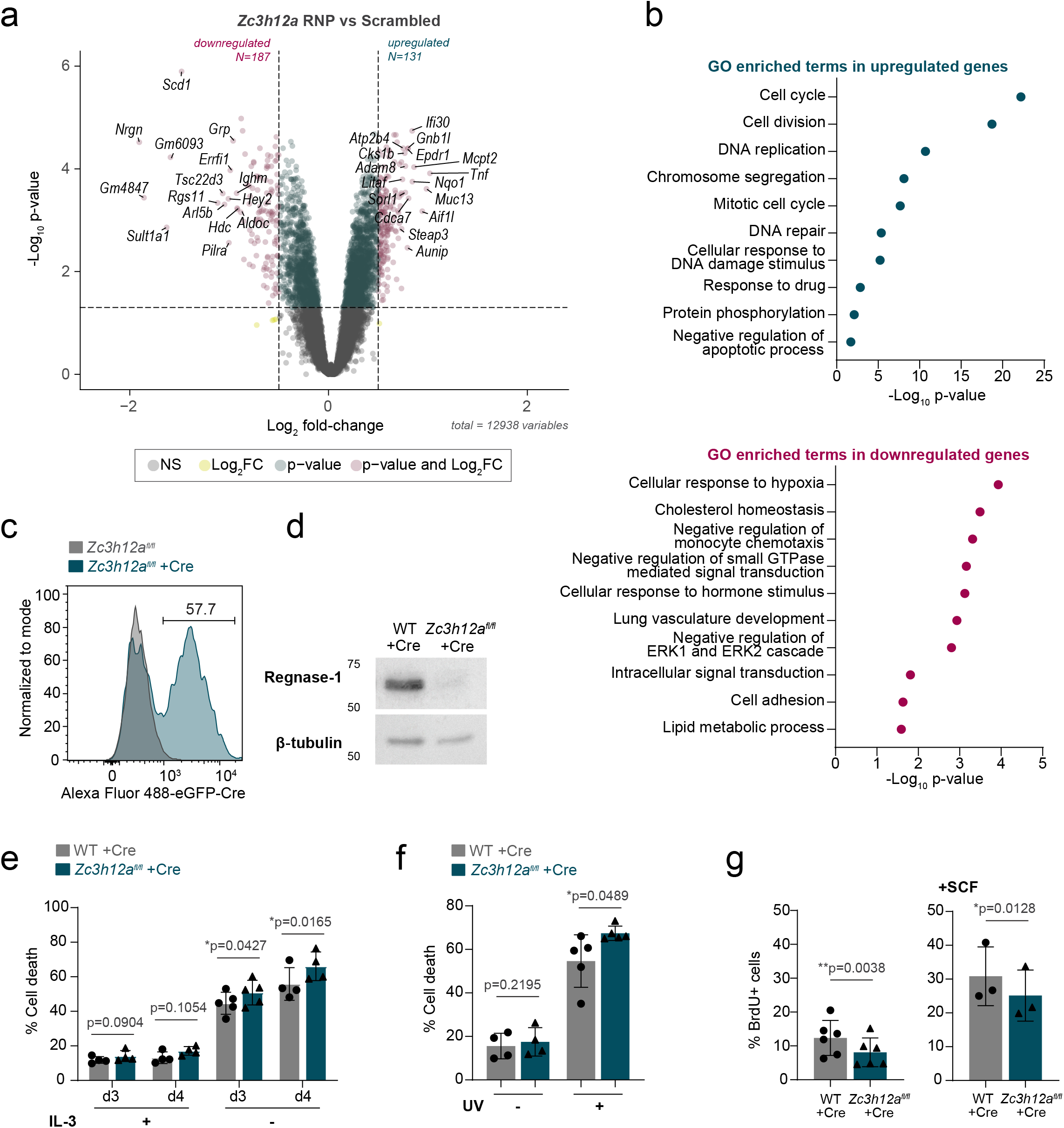
Loss of Regnase-1 induces widespread transcriptome changes in mast cells. **a)** Differential gene expression in BMMCs transfected with scrambled (control) or *Zc3h12a*-targeting Cas9-gRNA RNPs at resting state, measured by RNA-seq. **b)** Gene ontology analysis of enriched terms in upregulated and downregulated genes in **(a)**. **c)** Representative histogram of EGFP expression in WT and *Zc3h12a*^fl/fl^ BMMC transduced with EGFP-Cre recombinase-expressing lentivirus. Subsequent experiments were performed 4 to 7 days after transduction, gating on EGFP-positive cells. **d)** Expression of Regnase-1 in BMMCs differentiated from WT or *Zc3h12a*^fl/fl^ mice after 1 week of transduction with EGFP-Cre recombinase-expressing lentivirus, representative of N=4 independent experiments. **e)** LIVE/DEAD staining to measure cell viability after growing cells in medium with or without IL-3 for 72-96 hours. N=4 to 5 independent experiments. Mean ± SD. Paired t-test, two-tailed. **f)** Cell viability measured by LIVE/DEAD staining 72 h after UV treatment (200 J/m^2^). N=4 to 5 independent experiments. Mean ± SD. Paired t-test, two-tailed. **g)** BrdU staining to measure cell proliferation in normal culture medium containing IL-3 or supplemented with SCF. Mean ± SD. Paired t-test.

To verify this phenotype using an independent experimental approach, we took advantage of mast cells derived from the bone marrow of *Zc3h12a^fl/fl^*mice (30). Briefly, we transduced bone marrow cells with a lentiviral vector expressing an EGFP-Cre fusion protein (7) reaching at least 50% efficiency of transduction (**Figure 6c**). After sorting of EGFP^+^ cells, expression of Regnase-1 became undetectable by western blot (**Figure 6d**). Similar to cells treated with RNAi or Cas9 RNPs, *Zc3h12a^fl/fl^*-Cre mast cells showed reduced viability in different stress conditions, including the withdrawal of the survival factor IL-3 and short-term UV irradiation (**Figure 6e-f** and **Suppl. Figure 8c-d**). Proliferation was similarly reduced in these cells even in conditions that optimally sustain mast cell growth (addition of IL-3, with or without SCF) (**Figure 6g** and **Suppl. Figure 8e**). Overall, using a number of independent experimental approaches, we found that Regnase-1 is central to the maintenance of mast cell survival and basal proliferation, and that Regnase-1 deletion leads to widespread transcriptome changes associated to cell cycle defects.

## Discussion

In this study, we found that Regnase-1 and Regnase-3 contribute to the regulation of mast cell responses and homeostatic activities. Our data extend and complement previous studies showing negative regulation of inflammatory type 2 responses by Regnase-1 in other cell types. For instance, Regnase-1 was shown to modulate the activity of group 2 innate lymphoid cells, and the inhibition of Regnase-1 degradation in these cells led to attenuated pulmonary inflammation (31). Similarly, deletion of Regnase-1 in Th2 lymphocytes led to enhanced IL-5 production by these cells and lung inflammation (32). Compared to Regnase-1, current knowledge regarding the impact of Regnase-3 in modulating type-2 responses is more limited. Regnase-3 was primarily shown to modulate macrophage and dendritic cell responses, where it was shown to affect cytokine production, including TNF and IL-6 (22), although one of the primary role of Regnase-3 in macrophages was shown to be the regulation of Regnase-1 expression (17). We now showed that in mast cells the regulation of responses to IgE crosslinking requires the cooperative function of both Regnase-1 and Regnase-3. Among the genes differentially expressed upon Regnase-1 and −3 silencing, *Tnf* was found to be regulated by both proteins through their intrinsic enzymatic activities and by the direct targeting of the *Tnf* 3’UTR. An additive effect was clearly observed upon co-depletion of both Regnases, although differences in dosage or potency of RNase activity cannot be ruled out.

Post-transcriptional regulation of *Tnf* through the AU-rich elements (ARE) within its 3’UTR was previously described (33) and mostly attributed to the action of ARE-binding proteins, such as tristetrapolin (TTP) (34, 35). ARE-independent regulation through other regulatory elements was also reported, including a constitutive decay element (CDE) recognized by Roquin-1/2 (36, 37) and a new regulatory element (NRE) shown to be targeted by Regnase proteins, at least in reporter assays (38). Among these elements, removal of the ARE and NRE led to strong derepression of TNF expression and to embryonic lethality in mice (38, 39). In mast cells, TNF was previously reported to be regulated by TTP in an ARE-dependent manner in response to IL-4 (35), although not in response to stimulation with LPS (40), exemplifying context-dependent post-transcriptional regulation of this cytokine. Our data now provide evidence of the existence of an additional mechanism of TNF regulation exerted in mast cells by Regnase-1 and Regnase-3 in response to IgE stimulation. Regarding the specific effects of Regnase-1 or Regnase-3 on the *Tnf* 3’UTR, at least two scenarios can be envisioned. One possibility is that both proteins have exactly the same function, and any phenotypic difference is due to differences in their level of expression and subcellular localization. A second possibility is instead linked to differences in binding affinity and activity between the two proteins. These may be dependent on co-factors and post-translational modifications that can be at least in part signal- and cell-type specific. Importantly, we found that Regnase-3 is capable of stable interactions with Regnase-1, which may be important for their shared regulation of common inflammatory targets, such as *Tnf*. In previous studies, the interaction of Regnase-1 with other proteins, such as UPF1 and Roquin-1 (9, 24, 41), was shown to enhance its activity and specificity, while proteins involved in Regnase-1 post-translational modifications (MALT-1, IKKs/IRAK1, 14-3-3) influenced its stability and sub-cellular localization (8, 15, 23, 25). The protein interactors of Regnase-3, on the other hand, were so far largely unknown. Compared to Regnase-1, Regnase-3 shares homology in the PIN and zinc finger domains (17), but it also contains a long non-homologous C-terminal region that could serve as a hub for protein interactions that may endow Regnase-3 with unique roles.

In terms of gene expression, we found that basal expression of *Zc3h12a* and *Zc3h12c* were comparable in resting mast cells, but the extent of induction upon IgE-dependent activation was higher for *Zc3h12c* (∼30-folds after 2h, compared to a maximal induction for *Zc3h12a* of ∼3-folds). A similar, larger magnitude induction of Regnase-3 compared to Regnase-1 was also observed at protein level (∼4-fold *vs.* ∼2-fold, respectively). One possible interpretation of these findings is that Regnase-1 is primarily linked to the regulation of resting-state transcripts, while stimulus-induced inflammatory transcripts can be regulated by both proteins in a dose-dependent manner. Consistent with this possibility, we found that both Regnase-1 and −3 modulated the *Tnf* 3’UTR at a similar extent. However, the observation that these two proteins interact with each other but also with unique interactors raises the possibility that in intact cells the two proteins have indeed regulatory functions that are not fully redundant. Accordingly, HITS-CLIP analyses in transfected HEK293 cells showed an incomplete overlap in their target binding specificity (43). The fact that we found very limited co-localization of Regnase-1 and −3 even in overexpressing cells suggests that part of the incomplete target overlap may be linked to the physical segregation of these two proteins. The narrow influence of Regnase-3 on inflammatory gene expression may also be in part due to compensation by Regnase-1. Indeed, Regnase-3 negatively regulates Regnase-1 expression by destabilizing the *Zc3h12a* transcript through its RNase activity.

Consistent with a role of both Regnase-1 and Regnase-3 in co-regulating at least some stimulus-induced responses in mast cells, and with a predominant role of Regnase-1 on resting functions, concomitant knockdown of both proteins revealed mostly additive effects on inflammatory gene expression, suggesting cooperative and possibly redundant functions in restraining inflammation. However, in resting mast cells, only Regnase-1 was required for steady-state cell survival and proliferation. Regnase-3 knockdown did not exhibit an opposite phenotype, nor did the double knockdown resulted in any additive or antagonistic effects with respect to Regnase-1 knockdown alone. This implies that beyond dosage compensation, Regnase-1 may carry out unique roles in maintaining normal mast cell homeostasis.

Indeed, RNA-sequencing of mast cells lacking Regnase-1 revealed that many differentially expressed genes were associated with cell cycle and DNA replication, as well as with metabolic processes, including lipid and glucose metabolism. It is plausible that these phenotypes resulted at least in part from indirect effects of Regnase-1 deletion, since only few of the established direct Regnase-1 targets were dysregulated. For example, the increased cell death observed in mast cells lacking Regnase-1 is likely the result of cell cycle defects and replication stress, and of the overall poor cellular fitness observed in the absence of Regnase-1.

Overall, our study revealed that Regnase-1 and −3 constitute a network that cooperatively fine-tunes inflammatory gene expression during mast cell activation. Regnase-1 is additionally critical for mast cell survival and proliferation, while Regnase-3 is a direct interactor of Regnase-1 and is required to restrain Regnase-1 expression.

## Material and Methods

### Mice

All mice used in this study were on a C57BL/6 background, housed in a specific-pathogen free barrier facility under 12 h dark/light cycles, 20-24° C temperature and 50-65% humidity conditions. All animal studies were performed in accordance with Swiss Federal Veterinary Office guidelines and with approval from the Cantonal animal experimentation committee, Dipartimento della Sanità e della Socialità Cantone Ticino (authorization number TI10/19).

### Cell cultures

Bone marrow-derived mast cells (BMMCs) were *in vitro* differentiated from the bone marrow of 6-8-week old mice. Whole bone marrow cells were cultured in Iscove’s Modified Dulbecco’s Medium (Gibco) containing 10% heat-inactivated fetal bovine serum (Gibco) and IL-3 (recombinant, prepared in-house), and supplemented with 0.1 mM non-essential amino acids (Gibco), 2 mM L-alanyl-L-glutamine dipeptide (Gibco), 100 U/mL penicillin, 100 mg/mL streptomycin (Gibco), and 50 µM β-mercaptoethanol, as previously described (44, 45). After 4 weeks, ∼99% of the cells were FcεRIα^+^ c-Kit^+^ and were used for downstream analyses. Peritoneal mast cells (PMCs) were isolated *ex vivo* from mouse peritoneal cavity by intraperitoneal lavage followed by magnetic bead-based enrichment for c-Kit^+^ cells using anti-CD117 (c-Kit)-APC antibody (Biolegend) and APC microbeads (Miltenyi Biotec). Cells were expanded for at least 1 week in the presence of IL-3 and 30 ng/mL recombinant SCF (Peprotech) to obtain peritoneal cavity-derived mast cells (PCMCs). To generate mast cells lacking Regnase-1, bone marrow from *Zc3h12a*^fl/fl^ mice (30) were transduced with a lentiviral vector expressing the Cre recombinase fused to an EGFP reporter (7). Downstream analyses were performed by gating on EGFP^+^ cells (for flow cytometry-based assays) or by sorting for EGFP^+^ cells using BD FACSymphony S6 Cell Sorter (BD Biosciences) (for Western blot experiments). HEK293T cells were maintained in Dulbecco’s Modified Eagle Medium (Gibco) supplemented with 10% FBS (Gibco), 1 mM sodium pyruvate (Gibco), 100 U/mL penicillin, 100 mg/mL streptomycin (Gibco), and 50 µM β-mercaptoethanol.

### Mast cell activation

Mast cell cultures were stimulated with the following for the indicated time periods: 1 µg/ml IgE-anti DNP antibody (Sigma) and 0.2 µg/ml HSA-DNP (Sigma), 20 nM PMA (Sigma) and 2 µM ionomycin (Sigma), 10 ng/mL recombinant IL-33 (Biolegend), 1 µg/ml LPS (Invivogen), 10 ng/mL SCF (Peprotech).

### Reverse transcription-quantitative PCR (RT-qPCR)

Mast cells were lysed in TRI Reagent RT (Molecular Research Center) and total RNA was extracted using Direct-zol RNA microprep kit (Zymo Research) according to manufacturer’s protocol. cDNA was synthesized by reverse transcription using qScript cDNA SuperMix (Quanta Bioscience). Target genes were amplified using PerfeCTa SYBR Green FastMix (Quanta Bioscience) and QuantStudio 3 Real-Time PCR System (Thermo Fisher Scientific). All primer sequences used are listed in **Suppl. Table 5**. Data analysis was performed using the 2^-ΔΔCt^ or 2^-ΔCt^ method.

### Western blots

Mast cells were lysed in RIPA buffer (10 mM Tris-HCl pH 8.0, 1 mM EDTA, 1% Triton X-100, 0.1% sodium deoxycholate, 0.1% SDS, 140 mM NaCl) supplemented with protease inhibitor cocktail (Sigma) for protein detection. Protein concentration was quantified using Pierce BCA Protein Assay Kit (Thermo Fisher Scientific). Samples (50-100 μg) were run on SDS/polyacrylamide gels and blotted on a PVDF membrane using a wet transfer system. Membranes were blocked in 5% milk in TBST (5 mM Tris pH 7.3, 150 mM NaCl, 0.1% Tween-20) for 30 min at room temperature, then incubated with primary antibodies overnight at 4°C, followed by the corresponding HRP-conjugated secondary antibody for 1 h at room temperature. All antibodies used are listed in **Suppl. Table 5**. Chemiluminescence detection was performed using Clarity Western ECL Substrate (Bio-Rad) and Fusion FX7 EDGE Imaging System (Witec). Protein band intensity was quantified using ImageJ software version 1.53h (46).

### Surface marker and intracellular protein stainings

For surface marker staining, mast cells were stained using CD117 (c-Kit)-APC or APC/Cy7 and FcεRIα-PE (all from Biolegend) at 1:200 dilution for 20 min on ice. For all intracellular staining experiments, cells were washed with ice-cold PBS and stained with LIVE/DEAD™ Fixable Aqua or Blue Dead Cell Stain (Thermo Fisher Scientific) for 20 min at room temperature prior to cell fixation. For intracellular cytokine staining, cells were stimulated with different stimuli for 4 h. Brefeldin A (Sigma) was added in the last two hours of stimulation. Cells were then fixed in 4% paraformaldehyde for 10 min at room temperature, followed by permeabilization with 1% BSA-0.5% saponin solution in PBS, and staining with the following antibodies for 20 min on ice: TNF-α-PE/Cy7, IL-6-PE or APC (both from Biolegend) and IL-13-PE or eFluor 450 (eBioscience). For intracellular staining of HA-tagged proteins, cells were fixed and permeabilized using the eBioscience Foxp3 / Transcription Factor Staining Buffer Set (Thermo Fisher Scientific) according to manufacturer’s instructions. Cells were then stained with anti-HA.11 epitope tag (Biolegend) at 1:200 dilution for 1 h at room temperature, followed by anti-mouse IgG (H+L)-Alexa Fluor 647-conjugated secondary antibody at 1:500 dilution for 30 min at room temperature. All antibodies used are listed in **Suppl. Table 5**. Flow-cytometry data were collected using FACS Symphony A5 or LSRFortessa (BD Biosciences) and analyzed by FlowJo v10.6.0 (BD Biosciences).

### siRNA transfection

BMMCs were transfected with siRNA cocktails targeting *Zc3h12a* and/or *Zc3h12c* (Horizon Discovery/Dharmacon) using the 100 µl Neon Transfection System Kit (Thermo Fisher Scientific) according to manufacturer’s protocol. siGLO Green Transfection Indicator (Horizon Discovery/Dharmacon) was used as control. Briefly, cells were washed with PBS and resuspended in 100 µl of Buffer R. siRNAs (200 pmol) were then added to the cell suspension and cells were electroporated with one pulse at 1600V and 30 ms of width. Transfected cells were kept in antibiotic-free medium for 24 h. Downstream experiments were performed after 48-72 h. All siRNA sequences used are listed in **Suppl. Table 5**.

### Nanostring profiling

BMMCs transfected with siRNAs against *Zc3h12a* and/or *Zc3h12c* or siGLO control were stimulated with 1 μg/ml IgE–anti-DNP antibody and 0.2 μg/mL HSA-DNP antigen for 2 h, then lysed in TRI Reagent RT. Total RNA was extracted using Direct-zol RNA microprep kit and quantified using Qubit RNA High Sensitivity assay kit and Qubit 3 Fluorometer (Thermo Fisher Scientific). Purified RNA (50 ng) was hybridized to the nCounter Myeloid Innate Immunity Panel v2 codeset (NanoString Technologies) for 16 h at 65°C. Following hybridization, 30-35 µl of sample were added to the nCounter cartridge and analyzed using an nCounter SPRINT Profiler (NanoString Technologies) according to manufacturer’s instructions. Data analysis was carried out using the nSolver Analysis Software v4.0 (NanoString Technologies) and Omics Playground web-based platform (47). Genes with log_2_ fold-change >0.5 and <-0.5 and p-value ≤ 0.05 were considered as differentially expressed.

### CRISPR-Cas9 gene editing

BMMCs were transfected with Cas9-gRNA RNPs targeting *Zc3h12a* or *Zc3h12c* using the 10 µl Neon Transfection System Kit (Thermo Fisher Scientific) according to manufacturer’s protocol, and as previously described (7). To generate guide RNAs (gRNAs), equal amounts (400 pmol) of crRNA and tracrRNA were mixed with Nuclease Free Duplex buffer and annealed by boiling at 95°C then cooling down to room temperature. Three different crRNAs were selected for *Zc3h12a* and *Zc3h12c*. A non-targeting crRNA (scrambled) was used as control (all from Integrated DNA Technologies). Cas9-gRNA RNPs were prepared by incubating either 0.5 µl of each gRNA with 1.5 µl TrueCut Cas9 Protein v2 (5 µg/µl, Thermo Fisher Scientific) or 1 µl of each gRNA with 1.5 µl recombinant Cas9-NLS (5 µg/µl, in-house) for 20 min at room temperature. To increase transfection efficiency, Alt-R Electroporation enhancer (Integrated DNA Technologies) was added to the transfection mix. Cells were resuspended in 10 µl of Buffer R, and electroporated with the RNPs with one pulse at 1600V and 30 ms of width. Transfected cells were kept in antibiotic-free medium for 24 h. Downstream experiments were performed after 1 week. All oligonucleotides used are listed in **Suppl. Table 5**.

### Expression plasmids

Regnase-1 and Regnase-3 expression plasmids were generated using standard cloning techniques. The murine *Zc3h12a* coding sequence was amplified from pRetro-Xtight-Myc-GFP-Regnase-1 plasmid (24), while *Zc3h12c* coding sequence was amplified from cDNA obtained from BMMCs. These were either cloned into the pCDNA3 vector (Thermo Fisher Scientific) and pCDH-EF1α-T2A-copGFP (System Biosciences) for expression in HEK293T or tagged with FLAG-HA then cloned into the pUC57-mini vector (synthesized by Genscript) to be used as template for *in vitro* mRNA transcription. To generate the RNase-inactive mutants, residues D141 of Regnase-1 and D252 of Regnase-3 were mutated to asparigines using the QuikChange II XL Site-Directed Mutagenesis Kit (Agilent) following manufacturer’s protocol. To generate the luciferase reporter plasmids, the full-length 3’UTR of the *Tnf* and *Zc3h12a* genes was amplified from genomic DNA obtained from BMMCs and cloned downstream of the luciferase reporter gene in the pmirGLO Dual-Luciferase miRNA Target Expression Vector (Promega).

### *In vitro* mRNA transcription

To express Regnase-1 and −3, pUC57-mini plasmids harboring the WT or mutated coding sequence downstream of T7 promoter were first transcribed into mRNA using the HiScribe T7 ARCA mRNA Kit (New England BioLabs) according to manufacturer’s protocol. Briefly, the plasmids were linearized by digestion with SpeI (New England BioLabs), followed by purification using NucleoSpin Gel and PCR Clean-up kit (Macherey-Nagel). Linearized DNA templates (1 µg) were then assembled together with ARCA/NTP mix, T7 RNA polymerase mix (both included in the kit), and pseudo-UTP (Jena Bioscience) for the *in vitro* transcription (IVT) reaction. mRNA synthesis was completed by removing DNA template through DNase I treatment, followed by poly(A) tailing by adding Poly(A) polymerase and buffer (all included in the kit) to the reaction mix. All reactions were each done at 37°C for 30 min. Resulting mRNA products were purified using Monarch RNA Cleanup Kit (New England BioLabs), visualized on 1% TBE gel, and quantified using Nanodrop 2000. BMMCs were transfected with Regnase-1 or −3 (WT or RNase-inactive mutant D141N or D252N) IVT mRNA using the 10 µl Neon Transfection System Kit (Thermo Fisher Scientific) according to manufacturer’s protocol. IVT mRNA from pUC57-mini plasmid expressing ZsGreen alone was used as control. Briefly, cells were washed with PBS and resuspended in 10 µl of Buffer R. IVT mRNA (3 pmol) and 10U RNase inhibitor (Promega) were then added to the cell suspension. Cell electroporation was performed with one pulse at 1600V and 30 ms of width. Transfected cells were kept in antibiotic-free medium and downstream experiments were performed within 24 h.

### Luciferase reporter assays

HEK293T cells were transfected with 3 µg of pmirGLO plasmid containing the 3’UTR of *Zc3h12a* or *Tnf* (Addgene plasmid 207127) (7) and 1 µg of pCDNA3-Regnase-1 WT/D141N or pCDH-Regnase-3 WT/D252N or the corresponding empty control plasmids using polyethylenimine (PEI). After 48 h, luciferase activity was analyzed using the Dual-Luciferase Reporter Assay System (Promega) according to the manufacturer’s protocol and measured using GloMax Discover (Promega).

### mRNA stability assay

BMMCs were treated with 1 μg/ml IgE–anti-DNP (Sigma) for 30 min followed by stimulation with 0.2 μg/ml HSA-DNP (Sigma) for 30 min. Transcription was then blocked by incubating the cells with 10 μg/ml Actinomycin D (Sigma). Cells were harvested at different time points (15, 30, and 60 min), and RNA isolation and RT-qPCR were performed as mentioned above. mRNA decay rate (half-life, t^1/2^) was calculated by non-linear regression curve fitting (one phase decay) using Prism version 9 (GraphPad).

### Cell viability and apoptosis assays

Cell viability was measured by staining cells with LIVE/DEAD Fixable Aqua Dead Cell Stain or Blue Dead Cell Stain (Thermo Fisher Scientific) for 20 min at room temperature. Cell apoptosis was measured by co-staining cells with Annexin V and 7-AAD using the PE Annexin V Apoptosis Detection Kit I (BD Biosciences), according to manufacturer’s protocol.

### Cell proliferation and cell cycle analysis

Cell proliferation was measured by BrdU incorporation for 12 to 16 h at 37°C, followed by BrdU staining using the APC BrdU Flow Kit (BD Biosciences), according to manufacturer’s protocol. Cell cycle progression was analyzed by co-staining cells with 7-AAD.

### Cell degranulation assay

Mast cell degranulation assay was performed as previously described (44). Briefly, BMMCs were pre-incubated with 1 µg/ml IgE-anti DNP antibody overnight at 37°C. Cells were resuspended in Tyrode’s buffer (10 mM HEPES, 129 mM NaCl, 5 mM KCl, 15.05 mM BSA, 1.8 mM CaCl_2_, 2 mM MgCl_2_, 5 mM glucose) and stimulated with 0.2 μg/ml HSA-DNP for 1 h at 37°C. Cell supernatant was collected, while cell pellets were lysed in 0.5% Triton X-100 (Sigma) in Tyrode’s buffer. β-hexosaminidase substrate (4-nitrophenyl N-acetyl-β-D-glucosaminide, 4 mM, Sigma) was then added to the supernatant and lysate for 1 h at 37°C. The reaction was stopped with 0.2 M glycine (pH 10.7) and the absorbance was read at 405 nm. Percentage of cell degranulation was calculated as the ratio between the absorbance of the supernatant and the total absorbance (supernatant and cell lysates).

### RNA sequencing and bioinformatic analyses

BMMCs were transfected with *Zc3h12a* RNPs or scrambled RNPs as control (four independent biological replicates). After 1 week in culture, cells were lysed in TRI Reagent RT (Molecular Research Center) and RNA was extracted using Direct-zol RNA microprep kit (Zymo Research). After poly-A mRNA enrichment and Tecan Revelo mRNA library preparation, mRNA sequencing using Illumina Novaseq 6000 (2 x 50-bp reads) was outsourced to the Next Generation Sequencing Platform at the University of Bern (Switzerland). Read quality control was assessed using fastqc v.0.11.9 (http://www.bioinformatics.babraham.ac.uk/projects/fastqc/) and RSeQC v.4.0.0 (48), followed by read mapping to the reference genome Mus_musculus.GRCm39.107 using HiSat2 v.2.2.1 (49). Counts were generated and corrected for batch effects using featureCounts v.2.0.1 (50) and removeBatchEffect function in R (51), respectively. Differential expression was analyzed by combining three DE methods: DESeq2 (Wald), edgeR (QLF), and limma (trend), using the Omics Playground platform (47). Genes with log2 fold-change >0.5 and <-0.5 and p-value ≤ 0.05 were considered differentially expressed and included in the gene ontology (GO) analysis performed using DAVID Bioinformatics Resource v2023q2 (52, 53). The top 10 GO enriched terms in upregulated and downregulated genes were graphically represented (for upregulated genes, only categories with >10 genes were considered). Data visualization was performed with RStudio version 4.1.

Re-analysis of RNA-seq data from Li et al. (18) was performed using the DESeq2 package in R (54). Genes encoding for RBPs identified in more than one RBPome listed in the RBP2GO database (19) and with sum of counts in all samples ≥10 were included in the differential expression analysis. RBP genes with log2 fold-change >0.5 and <-0.5 and p-value ≤ 0.05 were considered differentially expressed and included in functional categorization analysis using PANTHER 17.0 (55). First, differentially expressed RBP genes broadly categorized under the GO term RNA metabolic process (GO:0016070) were identified. This was followed by more specific categorization using the following GO terms: regulation of RNA stability (GO:0043487), RNA modification (GO:0009451), RNA processing (GO:0006396), RNA localization (GO:0006403), and Translation (GO:0006412). Data visualization was performed with RStudio version 4.1. ATAC-seq data from Li et al. (18) were visualized using Integrative Genomics Viewer (IGV) 2.16.0 (56).

### Lentiviral preparation and cell transduction

To generate lentivirus, HEK293T cells were transfected with the lentiviral vector pScalps-EGFP-Cre recombinase (7) (Addgene plasmid 207132) together with the packaging vectors psPAX2 and pMD2.G (Addgene plasmids 12260 and 12259 by Didier Trono) using PEI. After 48 h, lentiviral particles in the supernatant were collected and concentrated using a PEG-8000 solution as previously described (57). Wild-type and *Zc3h12a*^fl/fl^ BMMCs were transduced with the lentivirus for at least 96 h. Flow cytometry-based analyses were done by gating on EGFP^+^ cells, while Western blot experiments were performed on EGFP^+^ cell population sorted using BD FACSymphony S6 Cell Sorter (BD Biosciences).

### Immunoprecipitation and enzymatic digestion

BMMCs (2×10^7^) were activated with 2 nM PMA and 200 nM ionomycin for 1 h. Cells were washed with ice-cold PBS and lysed in IP buffer (50 mM Tris pH 7.5, 150 mM NaCl, 1 mM EDTA, 0.25% NP-40, 1X protease inhibitor (Sigma), 1X phosphatase inhibitor (Sigma)). Lysates were pre-cleared by incubating with 20 µL Dynabeads Protein G (30 mg/ml, Invitrogen) at 4°C with rotation for 2 h. 1 mg of pre-cleared lysates was incubated with 50 µL Dynabeads Protein G (30 mg/ml) and 5 µg Regnase-1 (R&D systems) or Regnase-3 (Helmholtz Munich Monoclonal Antibody Core Facility) antibody at 4°C with rotation for 16 h. Mouse IgG (Sigma) or rat IgG (Thermo Fisher Scientific) antibodies were used as isotype controls, respectively. Immunoprecipitated proteins were pulled down using DynaMag-2 Magnet (Thermo Fisher Scientific) and washed 2 times with IP buffer with 0.1% NP-40, followed by 2 washes with detergent-free wash buffer (50 mM Tris pH 7.5, 150 mM NaCl, 1X protease inhibitor, 1X phosphatase inhibitor).

On-bead digestion of immunoprecipitated proteins was performed as follows. Beads were resuspended in 8M urea, 50 mM ammonium bicarbonate buffer. Proteins were then reduced with 10 mM dithiothreitol for 60 minutes at 37°C and alkylated with 50 mM iodoacetamide for 30 min at room temperature. Digestion was carried out in 8M urea, 50 mM ammonium bicarbonate buffer for 2 hours at 37°C with 1 µg of Lys-C (FUJIFILM Wako Chemicals), after which the digestion buffer was diluted to final 2M urea with 50 mM ammonium bicarbonate. 1 µg of trypsin (Promega) was added for overnight digestion at 37°C. All steps were performed in agitation to avoid beads precipitation. Digestion was stopped by adding acetonitrile to 2% and trifluoroacetic acid to 0.3% and the beads were collected with DynaMag-2 Magnet (Thermo Fisher Scientific). Digested peptides were purified by loading the supernatant into C18 StageTips (58), and eluted with 80% acetonitrile, 0.5% acetic acid. Finally, the elution buffer was evaporated by vacuum centrifugation and purified peptides were resolved in 2% acetonitrile, 0.5% acetic acid, 0.1% trifluoroacetic acid for single-shot LC-MS/MS measurements.

### LC-MS/MS

Peptides were separated on an EASY-nLC 1200 HPLC system (Thermo Fisher Scientific) coupled online via a nanoelectrospray source (Thermo Fisher Scientific) to a Q Exactive HF mass spectrometer (Thermo Fisher Scientific). Peptides were loaded in buffer A (0.1% formic acid) into a 75 µm inner diameter, 50 cm length column, packed in-house with ReproSil-Pur 120 C18-AQ 1.9 µm resin (Dr. Maisch HPLC GmbH), and eluted over a 150-min linear gradient of 5 to 30% buffer B (80% acetonitrile, 0.1% formic acid) at a flow rate of 250 nl/min. The Xcalibur software (Thermo Scientific) operated the Q Exactive HF in a data-dependent mode with a survey scan range of 300-1,650 m/z, resolution of 60,000 at 200 m/z, maximum injection time of 20 ms and AGC target of 3e6. The top-10 most abundant ions with charge 2 to 5 were isolated with a 1.8 m/z isolation window and fragmented by higher-energy collisional dissociation (HCD) at a normalized collision energy of 27. MS/MS spectra were acquired with a resolution of 15,000 at 200 m/z, maximum injection time of 55 ms and AGC target of 1e5. Dynamic exclusion was set to 30 s to avoid repeated sequencing.

### LC-MS/MS data analysis

MS raw files were processed using the MaxQuant software v.1.6.7.0 (59). Peptides and proteins were identified with a 0.01 false discovery rate (FDR) by employing the integrated Andromeda search engine (60) to search spectra against the Mouse UniProt database (July 2019) and a common contaminants database (247 entries). Enzyme specificity was set as “Trypsin/P” with a maximum of 2 missed cleavages and minimum length of 7 amino acids. N-terminal protein acetylation and methionine oxidation were set as variable modifications, and cysteine carbamidomethylation as a fixed modification. Match between runs was enabled to transfer identifications across samples based on mass and normalized retention times, with a matching time window of 0.7 min and an alignment time window of 20 min. Label-free protein quantification (LFQ) was performed with the MaxLFQ algorithm (61) with a minimum required peptide ratio count of 1. Data analysis was performed using the Perseus software v.1.6.2.3 (62). Data were pre-processed by removing proteins only identified by site, reverse hits, and potential contaminants. After log2 transformation of LFQ intensities, biological replicates of each experimental condition were grouped and proteins were filtered for a minimum of 3 valid values in at least one group. Missing data points were then replaced by imputation from a normal distribution with 0.3 width and 1.8 downshift, and a two-sided two-samples t-test (0.05 FDR, 250 randomizations) was used to identify significant changes of protein intensity between each immunoprecipitation experiment and its corresponding isotype control.

### Regnase-1 and −3 co-transfection in HEK293T cells

HEK293T cells were transfected with pCDNA3-Regnase-1 and pCDNA3-Regnase-3 plasmids (10 µg each) using polyethylenimine (PEI). After 48 h, cells were harvested for immunoprecipitation experiments.

### Immunofluorescence

Cells were fixed with 4% paraformaldehyde in PBS for 10 min, then cytospun on a coverslip. Cells were permeabilized with 0.2% Triton X-100 in PBS for 15 min, and blocked with 5% BSA in PBS for 30 min. Cells were then stained with anti-Regnase-1 (R&D systems) or anti-Regnase-3 (Helmholtz Munich Monoclonal Antibody Core Facility) at 1:100 dilution for 1.5 h, followed by anti-mouse IgG (H+L)-Alexa Fluor 568- or anti-rat IgG (H+L)-Alexa Fluor 647-conjugated secondary antibody (Thermo Fisher Scientific) at 1:500 dilution for 30 min. All incubations were done at room temperature. Stained cells were washed three times with PBS and once with water, followed by nuclei counterstaining and mounting on microscopy slides using Vectashield (Vector Laboratories) with 4′,6-diamidino-2-phenylindole (DAPI). All antibodies used are listed in **Suppl. Table 5**. Confocal microscopy imaging was performed using Leica Stellaris SP8 confocal laser scanning microscope with Leica HCX PL APO 40×1.30 oil or 63×1.40 oil objectives (Leica Microsystems). Image acquisition and processing were done using Leica Application Suite X (Leica Microsystems) and ImageJ version 1.53h (46) softwares.

### Data analysis

All data representation and statistical analysis were performed using GraphPad Prism v9. All source data for all figures are provided in the **Source Data File**.

## Data availability statement

All raw data are either contained within this manuscript or are deposited in Gene Expression Omnibus (GEO) with accession number GSE240210.

## Author contributions

Conceptualization: MB, CL, SM. Data generation: MB, CL, MP, EW. Data analysis: MB, CL, SMoro, MP.

Paper writing: MB, SM with contributions from all the authors. Supervision: SM.

Funding acquisition: SM, VH.

## Acknowledgements

Work on RNA-binding proteins in the SM lab was funded by the Swiss National Science Foundation grant number 310030L_189352 and the Aldo and Cele Daccò Foundation (to SM). The authors would like to thank Pamela Nicholson (Next Generation Sequencing Platform, University of Bern) for RNA-seq.

